# A genomic tool to tackle cryptic diversity demonstrates the potential for off-target use of GT-seq panels

**DOI:** 10.64898/2026.06.09.731139

**Authors:** Amanda S. Ackiss, Mark R. Vinson, Ann J. Ropp, Kristen M. Gruenthal, Trevor J. Krabbenhoft, Joseph V. Siegel, Wendylee Stott, Daniel L. Yule, Wesley A. Larson

**Author notes:** retired (Mark R. Vinson).

## Abstract

A comprehensive understanding of life history is vital to successful species conservation and management. When different life history stages are accompanied by considerable morphological or cryptic variation, such as the egg and larval phases exhibited by most fishes, genomic tools are essential for identifying species so that early-life ecology questions can be studied. Genotyping-in-thousands by sequencing (GT-seq) has recently emerged as a targeted and efficient approach for species identification. We leveraged existing genomic and transcriptomic data to develop a GT-seq panel capable of differentiating the members of the *Coregonus artedi* complex, a radiation of salmonids in the Laurentian Great Lakes whose members are indistinguishable with mitochondrial DNA barcoding loci and are the focus of bi-national conservation initiatives. Our panel of 494 loci was able to assign fishes in the *C. artedi* complex to species and lake. We examined cross-amplification in other coregonines with overlapping distributions and found that congeneric Lake Whitefish (*C. clupeaformis*) cross-amplified at 94% of loci and confamilial Round and Pygmy Whitefish (*Prosopium* spp.) cross-amplified at 42% and 38% of loci, respectively. We adapted bioinformatic probes to account for *Prosopium*-specific variants including 22 new SNPs and developed a whitelist of 428 SNPs capable of distinguishing these whitefishes. Finally, we demonstrated performance by identifying 3,066 coregonine larvae and juveniles collected in spring 2019-2021 from Lake Superior. These results hold promise for future insights into the species-specific ecology of early life coregonines and demonstrate the flexibility of GT-seq panels, which may cross-amplify hundreds of informative genome-wide loci in related taxa.

## Introduction

Cryptic diversity poses many challenges to successful species conservation and management (Hending, 2025). The ability to identify species throughout their life cycle is vital to understanding their ecology and biology, but if a species is elusive (Claridge et al. 2004) or its phenotypes are visually indistinguishable (Bickford et al., 2007), fundamental gaps in knowledge can arise with potentially dire consequences (Geller, 1999; Schönrogge et al., 2002; Nguyen et al., 2019). In many species, cryptic diversity may be especially prevalent in early life stages such as propagules and larvae (Pfenninger et al. 2007; Collin et al., 2020; Hoffman et al., 2021). In others, cryptic diversity occurs in adults where morphological differences may be negligible (Hebert et al., 2004; Vega-Sánchez et al., 2024). In both cases, molecular markers have become one of the most reliable means for detecting cryptic diversity (Hebert et al., 2003; Stoeckle 2003; Bickford et al., 2007), and high-throughput methods for species identification at both individual and community levels are becoming increasingly common with decreasing sequencing costs (Kimmerling et al., 2018; Porter & Hajibabaei, 2018; Miya et al., 2020; Hebert et al., 2025).

Recent evolutionary divergence has been identified as a potential mechanism causing cryptic diversity (Chenuil et al., 2018; Struck et al., 2018; Cahill et al., 2024), and it can complicate genetic species identification even in the absence of morphological ambiguity. Success rates for genetic identification of recently diverged species with standard mitochondrial, chloroplast, and nuclear DNA barcoding markers have been shown to be significantly lower than for evolutionarily older species due to factors such as incomplete lineage sorting (van Velzen et al., 2012). In such cases, increasing marker density across the genome improves discrimination of distinct evolutionary units. For example, two of the four subspecies in the North American yellow-rumped warbler (*Setophaga coronata*) species complex could not be reliability distinguished with genetic methods until high throughput, reduced-representation or whole genome sequencing was used (Toews et al., 2016; Szarmach et al., 2021). Similarly, restriction site-associated DNA sequencing (RAD-seq) was able to completely resolve reciprocal monophyly across 16 recently diverged Lake Victoria cichlids for the first time (Wagner et al., 2013).

Reduced representation and whole genome sequencing are powerful methods to address the challenges of characterizing cryptic or recently diverged populations and species and are more accessible than ever for non-model organisms (Ekblom & Galindo, 2011; Ellegren, 2014); however, more cost-effective approaches for genetic identification are needed when repeated or large numbers of samples are required to address specific questions. Over the last decade, targeted amplicon panels such as genotyping-in-thousands by sequencing (GT-seq; Campbell et al., 2015) have emerged as a practical approach to combining the power of genome-wide loci and the need for fast, cost-effective genotyping (Meek & Larson, 2019). Developed from reduced representation sequencing (Bootsma et al., 2020; Euclide et al., 2022; Hayward et al., 2022, Homola et al. 2025) and whole genome sequencing (Beemelmanns et al., 2025) datasets, amplicon panels can be designed to target both neutral and adaptive loci (Garrett et al., 2024; Schmidt et al., 2020) enabling a broad range of population and demographic inference. Since the amplification target is a sequence rather than specific single nucleotide polymorphisms (SNPs), which are the target of widely used, cost-effective assays such as TaqMan® and high-density SNP chips (Francisco et al., 2005; Gunderson et al., 2006; Groenen et al., 2011), amplicons that contain multiple SNPs confer the additional benefit of calling microhaplotypes (Baetscher et al., 2018; Baetscher et al., 2023). As a growing number of amplicon panels are developed for genetic monitoring, there is potential for leveraging existing panels for off-target species through cross-amplification as was commonly done during the age of microsatellites (Moore et al., 1991; Schlotteröer et al., 1991; Primmer et al., 1996; Wilson et al., 2004).

The *Coregonus artedi* species complex, a radiation of salmonids in the Laurentian Great Lakes of North America known as ciscoes, embodies the challenges of both cryptic diversity and recent evolutionary divergence. The *Coregoninae* subfamily underwent repeated diversification events in cold-water habitats across the northern hemisphere after the Last Glacial Maximum (Hudson et al., 2007), and the short evolutionary timescale across which this diversification occurred has led to uninformative and often conflicting results when using classical genetic and morphological data to infer species relationships (Nikolsky & Reshetnikov, 1970; Reed et al., 1998; Turgeon et al., 1999; Turgeon & Bernatchez, 2003), a situation referred to as the “coregonid problem” (Svardson 1949; Stott & Todd, 2007; Mee et al., 2015). Additionally in the Great Lakes, many deepwater species in this complex can be difficult to visually distinguish (Koelz, 1929), and there are no morphological characters that can reliably identify the eggs or larvae (Auer, 1982; George et al., 2018) rendering critical gaps in our understanding of species-specific early life history. Recent studies on Great Lakes ciscoes employing RAD-seq (Ackiss et al., 2020; Lachance et al., 2021), RNA-seq (Bernal et al., 2022), and whole genome sequencing (Backenstose et al., 2024) have enabled reliable and accurate species identification, particularly among deepwater species, for the first time.

A century of severe declines in the diversity of Great Lakes ciscoes (Eshenroder et al., 2016) has led to the development of a bi-national framework to support coregonine restoration (Bunnell et al., 2023). The framework recognizes a need to resolve species relationships with genetics and augments previously identified research needs for improving our understanding of coregonine ecology, including species-specific larval emergence timing, dispersal, and duration and the influence of environmental and invasive species interactions on recruitment variability (Zimmerman & Krueger, 2009). We leveraged recent genomic and transcriptomic datasets to develop a GT-seq panel capable of discriminating ciscoes that will allow scientists to cost-effectively address these questions at a basin-wide scale. Since early life stages (larvae, eggs) could be confused with those of sister species, including Lake Whitefish (*C. clupeaformis*), Round Whitefish (*Prosopium cylindraceum*), and Pygmy Whitefish (*P. coulterii*), we also tested the cross-amplification of panel loci and successfully modified a subset of genotyping probes for whitefish species identification. Finally, we used the panel to identify 3,066 larval and age-1 coregonines sampled in Lake Superior in 2019-2021 enabling the first large scale examination of early life diversity and distribution in Great Lakes ciscoes. Our results lay the foundation for a myriad of insights into coregonine early life history that can inform their conservation and restoration and demonstrate the ease with which GT-seq panels can be adapted for off-target use.

## Methods

### Development of an amplicon panel from RAD-seq data

Candidate loci for an amplicon panel for genetic stock identification (GSI) of Great Lakes ciscoes were identified from RAD-seq data from sampled adults representing the three major extant species in the *Coregonus artedi* complex (*C. artedi*, *C. hoyi*, *C. kiyi*). Two other species, *C. zenithicus* and *C. nigripinnis*, have uncertain status Lake Superior and may potentially be extinct (Eshenroder et al. 2016). Fishes field identified as *C. zenithicus* and *C. nigripinnis* are caught extremely rarely, and RAD-seq data from these fishes lend uncertainty as to whether those field identifications are correct (Ackiss et al. 2020), therefore, *C. zenithicus* and *C. nigripinnis* were not included in this analysis. Lake-specific representatives of each species were also included (Superior, Michigan, Huron and Ontario, Figure 1; Table 1). DNA from samples was extracted using Qiagen DNeasy Blood & Tissue kits, quantified using a Quant-it™ PicoGreen^®^ dsDNA Assay (Invitrogen, Waltham, MA), and normalized to 200 ng for bestRAD library preparation which followed the same protocols outlined by Ackiss et al. (2020). Libraries were sequenced through Novogene (Sacramento, CA) using paired-end (PE) 150 technology on the Illumina Novaseq platform.

**Figure 1.**
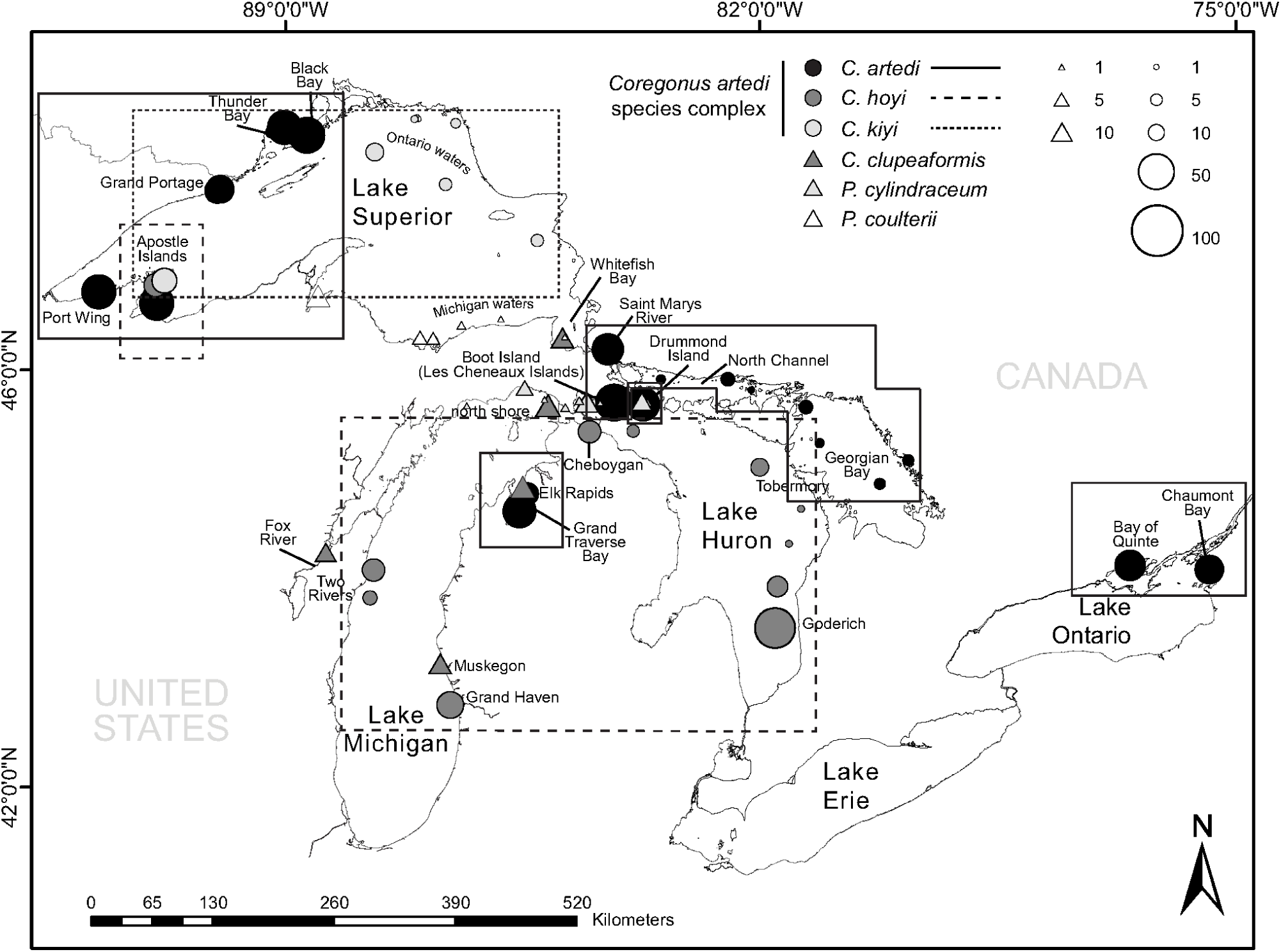
Map of collection sites across the Great Lakes for samples used in GT-seq panel development (circles) and cross-amplification testing (triangles). Boxes indicate reporting groups for Cisco *C. artedi* (solid lines), Bloater *C. hoyi* (dashed lines), and Kiyi *C. kiyi* (dotted line). Sizes of collection site markers scaled to number of samples (see Table 1 for collection and reporting group details).

**Table 1.**
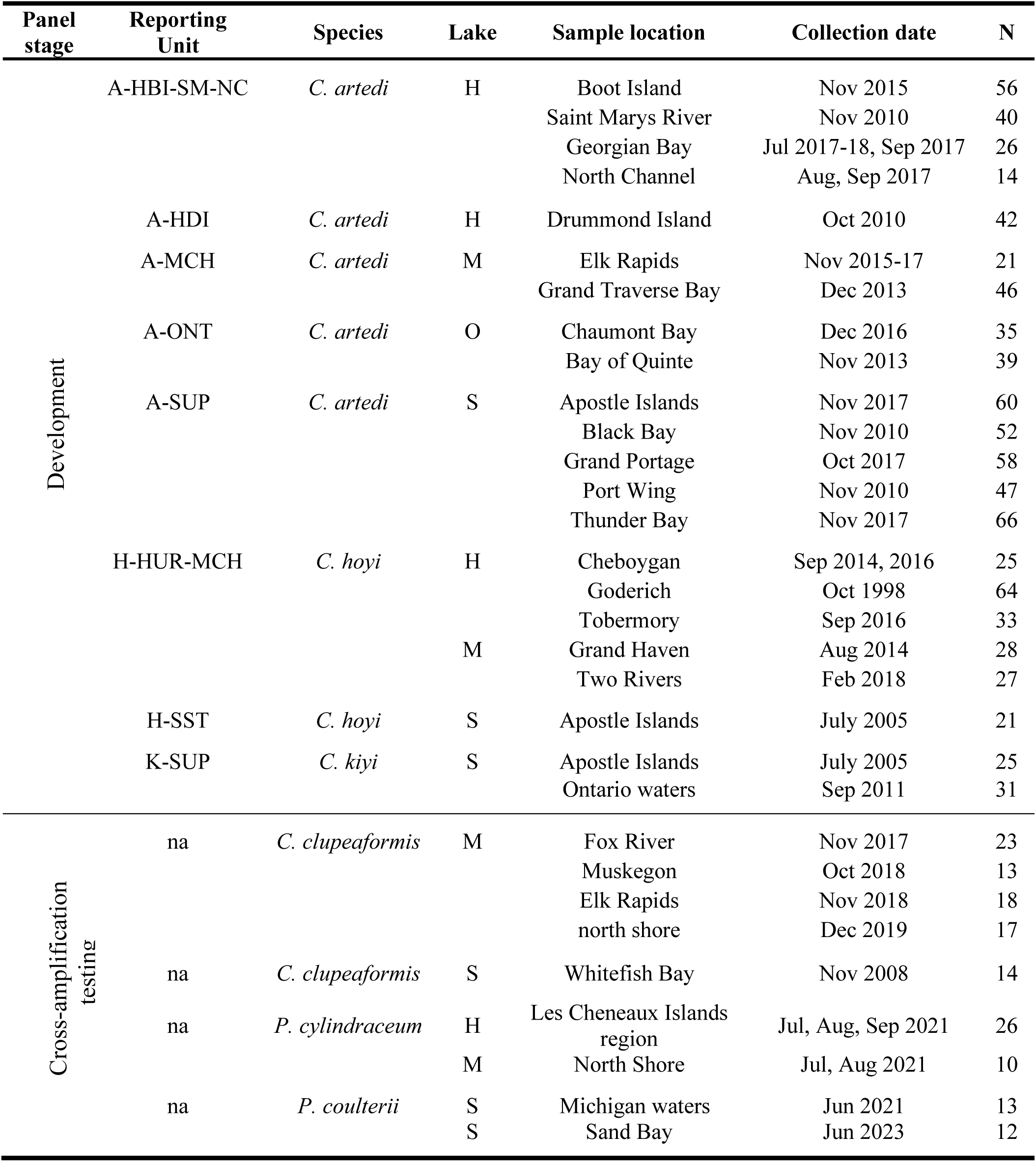
Collections used for GT-seq panel development (identification and simulated performance of candidate loci from RAD-seq data) and cross-amplification testing on sister species and genera. Lake codes are Huron (H), Ontario (O), Michigan (M), Superior (S), **N** is the number of genotyped samples from each population, **na** = not applicable.

| Panel stage | Reporting Unit | Species | Lake | Sample location | Collection date | N |
| --- | --- | --- | --- | --- | --- | --- |
| Development | A-HBI-SM-NC | *C. artedi* | H | Boot Island | Nov 2015 | 56 |
|  |  |  |  | Saint Marys River | Nov 2010 | 40 |
|  |  |  |  | Georgian Bay | Jul 2017-18, Sep 2017 | 26 |
|  |  |  |  | North Channel | Aug, Sep 2017 | 14 |
|  | A-HDI | *C. artedi* | H | Drummond Island | Oct 2010 | 42 |
|  | A-MCH | *C. artedi* | M | Elk Rapids | Nov 2015-17 | 21 |
|  |  |  |  | Grand Traverse Bay | Dec 2013 | 46 |
|  | A-ONT | *C. artedi* | O | Chaumont Bay | Dec 2016 | 35 |
|  |  |  |  | Bay of Quinte | Nov 2013 | 39 |
|  | A-SUP | *C. artedi* | S | Apostle Islands | Nov 2017 | 60 |
|  |  |  |  | Black Bay | Nov 2010 | 52 |
|  |  |  |  | Grand Portage | Oct 2017 | 58 |
|  |  |  |  | Port Wing | Nov 2010 | 47 |
|  |  |  |  | Thunder Bay | Nov 2017 | 66 |
|  | H-HUR-MCH | *C. hoyi* | H | Cheboygan | Sep 2014, 2016 | 25 |
|  |  |  |  | Goderich | Oct 1998 | 64 |
|  |  |  |  | Tobermory | Sep 2016 | 33 |
|  |  |  | M | Grand Haven | Aug 2014 | 28 |
|  |  |  |  | Two Rivers | Feb 2018 | 27 |
|  | H-SST | *C. hoyi* | S | Apostle Islands | July 2005 | 21 |
|  | K-SUP | *C. kiyi* | S | Apostle Islands | July 2005 | 25 |
|  |  |  |  | Ontario waters | Sep 2011 | 31 |
| Cross-amplification testing | na | *C. clupeaformis* | M | Fox River | Nov 2017 | 23 |
|  |  |  |  | Muskegon | Oct 2018 | 13 |
|  |  |  |  | Elk Rapids | Nov 2018 | 18 |
|  |  |  |  | north shore | Dec 2019 | 17 |
|  | na | *C. clupeaformis* | S | Whitefish Bay | Nov 2008 | 14 |
|  | na | *P. cylindraceum* | H | Les Cheneaux Islands region | Jul, Aug, Sep 2021 | 26 |
|  |  |  | M | North Shore | Jul, Aug 2021 | 10 |
|  | na | *P. coulterii* | S | Michigan waters | Jun 2021 | 13 |
|  |  |  | S | Sand Bay | Jun 2023 | 12 |

Reads were processed in STACKS to identify variant loci using the same parameters for the subprograms *process_radtags*, *ustacks*, *cstacks*, *gstacks*, and *populations* described in Ackiss et al. (2020). The *de novo* catalog generated in *cstacks* was created with five individuals from each unique species and lake. Iterative SNP filtering was performed with vcftools v0.1.15 (Danecek et al., 2011) and included (1) removing loci with a minor allele count less than three, (2) removing loci genotyped in fewer than 30% of individuals, (2) removing individuals missing more than 50% of loci, and (4) removing remaining loci genotyped in fewer than 70% of individuals. Putatively paralogous loci were identified with the program HDPlot (McKinney et al., 2017) and filtered using cutoffs of heterozygosity >0.55 and a read ratio deviation >5 and <–5. In order to identify candidate loci for genetic stock identification (GSI) and the reporting groups to test them, G-statistics (analogues of F-statistics based on Nei, 1987) including differentiation (*Gʹ*_ST_), inbreeding coefficients (*G*_IS_), and observed and expected heterozygosity (*H*_o_/*H*_e_), were calculated for each locus and collection in Genodive v3.0 (Meirmans, 2020).

### Simulated performance of candidate panels for genetic stock identification

Reporting groups for assessing candidate panel performance were created by grouping individuals from unique collection sites and species with pairwise *Gʹ*_ST_ >0.015. Self-assignment of individuals to reporting groups was assessed using a leave-one-out procedure with the *self_assign* function in the R package *rubias* (Moran & Anderson, 2019). Candidate panels for genetic stock identification of 600 RAD-seq loci each were designed with different combinations of high *Gʹ*_ST_ loci and high diversity loci generated from basin-wide and Lake Superior only datasets as both SNP and haplotype test datasets (Supplemental File 1, Supp. Table 1), following similar protocols outlined by Bootsma et al. (2020) and Euclide et al. (2022). The performance of candidate panels was simulated using the *assess_reference_loo* function in *rubias* (mix size = 200, MCMC simulations = 1000).

### GT-seq panel development

Primers for candidate loci were designed in Geneious Prime v.2019.2 using locus consensus sequences pulled from the STACKS catalog (catalog.fa) and a vcf file containing all polymorphic sites on a target locus. Primer design parameters included an optimal product size of 100bp (min 50bp, max 150bp) and optimal probes with a size of 20bp (min 18bp, max 25bp), Tm of 60℃ (min 57℃, max 63℃), GC content of 50% (min 40%, max 60%) and homopolymers (poly-X) no longer than three bps. The Test Primers tool in Geneious was used to check whether primers would bind non-target loci in the pool of consensus sequences. Primers binding non-target fragments were redesigned, and if redesign did not fix non-target binding or was not possible, the locus was dropped. Primer design occurred iteratively; as loci were dropped due to lack of optimal priming sites or primers that bound non-target fragments, the locus with the next highest *Gʹ*_ST_ value was selected for primer design until a total of 600 primer sets was reached. In addition to RAD-seq loci, primers were also designed for 45 loci from transcriptomic data that contained a polymorphic site exhibiting high differentiation among *C. artedi*, *C. hoyi* and *C. kiyi* in Lake Superior (Bernal et al., 2022). Primer design for transcriptomic data followed the same protocol as above, using cDNA sequences annotated with a vcf file containing information on target variant location.

GT-seq panel optimization was performed following the methods described by Bootsma et al. (2020) modified from the original methods of Campbell et al. (2015). Briefly, samples were amplified at panel loci, barcoded with i5/i7 indices, normalized, and pooled into master libraries for sequencing with PE 150 chemistry on the Illumina sequencing platform. Detailed panel amplification parameters can be found in supplemental information (Supplement File 2). Locus genotyping and evaluation was performed using the GTscore pipeline (https://github.com/gjmckinney/GTscore), which uses a Perl script and probe file designed to quickly identify loci and variants from a list of forward primer sequences and 20 bp probe sequences that identify target alleles. The original 645 GT-seq panel loci were iteratively reduced over two rounds of test amplification. Over-represented loci were identified and removed after each round of test amplification and sequencing. The STACKS catalog sequences for the final subset of panel loci were mapped to the *C. artedi* genome (GCA_039881085.1; Backenstose et al., 2024) with BWA fastmap (Li, 2013) to characterize coverage across the genome. Loci composed of super-maximal exact matches with 2 or fewer mismatches to chromosomes were visualized with karyoploteR (Gel & Serra, 2017).

### Genotyping reference individuals and testing cross-amplification

We selected a subset of ciscoes to serve as reference individuals for genetic assignments and genotyped them with the final panel of GT-seq loci. Reference individuals were comprised of adult fishes not used in the RADseq analysis for panel development that were morphologically identified as Cisco *C. artedi* (n=30), Bloater *C. hoyi* (n=49), and Kiyi *C. kiyi* (n=50) and collected from Lake Superior between 2014-2021 (ScienceBase, https://doi.org/10.5066/P13W68YB). DNA extraction of tissue from reference individuals was performed with Qiagen® DNeasy Blood & Tissue kits following manufacturer’s protocols with a final elution in 200ul Tris low-EDTA (TLE, 10mM Tris-HCl, 0.1 mM EDTA) buffer.

To explore cross-amplification of panel loci in other coregonines that can be difficult to distinguish from ciscoes in early life stages, we tested panel amplification on the other three coregonines present in the Great Lakes: Lake Whitefish (*Coregonus clupeaformis*), Round Whitefish (*Prosopium cylindraceum*), and Pygmy Whitefish (*P. coulterii*). Lake Whitefish samples included adults collected from four locations in Lake Michigan between 2017-2019 (n=72) and one location in Lake Superior in 2008 (n=14). *P. cylindraceum* samples included adults collected in Lake Michigan and Lake Huron in 2021 (n=36). *P. coulterii*, which are only present in Lake Superior, were also collected in 2021 and 2023 (n=25). Collection locations of outgroup samples are shown in Fig. 1, and sample metadata are archived on ScienceBase (https://doi.org/10.5066/P13W68YB). DNA extraction of outgroup sample tissues was performed with Qiagen® DNeasy Blood & Tissue kits as described for reference ciscoes, except for the Lake Whitefish samples from Lake Michigan, which were extracted using a 10% Chelex 100 (200-400 mesh; Biorad, Hercules, CA) slurry as described in Bootsma et al. (2020).

GT-seq libraries for all reference individuals were prepared with the final panel following the protocol described in Supplement File 2. Master library pools were PE150 sequenced with v2 chemistry on an Illumina MiSeq at the US Geological Survey’s Great Lakes Science Center (Ann Arbor, Michigan, USA). Genotypes were called from forward reads using a probe file in GTscore.

### Quantifying cross-amplification success and modifying probes for outgroup assignment

We quantified the success of cross amplification of the GT-seq panel on congener and confamilial whitefishes with three methods. First, we evaluated the number of loci and SNPs that were genotyped for Great Lakes whitefishes in GTscore with a file used to call SNPs via 20 bp in silico probes for identifying ciscoes. Each whitefish species was run separately, and loci and SNPs that genotyped in >50% of samples were counted. GT-seq loci that cross-amplified were annotated on the 38 chromosomes of the *C. artedi* genome with karyoploteR (Gel & Serra, 2017). We mapped outgroup reads to the FASTA catalog of loci in the panel with BWA mem (Li, 2013) and examined loci that cross-amplified in whitefishes in IGV. Our intent was to modify existing panel probes to expand their capacity for whitefish identification rather than to create entirely new probes which could introduce variability to our panel and potentially reduce accuracy for identification of ciscoes. If existing SNP probes overlapped a variant that was present in a whitefish species, the probe was modified to account for and/or call the SNP. Probes that overlapped multiple whitefish SNPs were not redesigned. The newly modified probe file was used to genotype references with GTscore, and genotyping success was examined in the dartR package in R (Gruber et al., 2018; Mijangos et al., 2022). The number of panel loci that cross-amplified in Round and Pygmy Whitefishes (*Prosopium* spp.) was much lower than the number that cross amplified in sister species Lake Whitefish (*Coregonus*), therefore, we tested the genetic assignment power of 1) a whitelist of the panel loci that were polymorphic in a dataset comprised of all whitefishes and 2) a whitelist of the panel loci that were polymorphic in a dataset comprised of only *Prosopium* whitefishes. For both, individuals that were missing >25% of the data were dropped and assignment accuracy was tested with the assign.MC function in the R package *assignPOP* (Chen et al., 2018). Training data included proportions of 0.5, 0.7, and 0.9 for individuals and 0.1, 0.25, 0.5, and 1 for loci over 30 iterations with the support vector machine (svm) predictive model.

Second, we mapped forward reads from Great Lakes whitefishes to the *C. artedi* genome to quantify locus amplification and number of variant sites sampled within species. Adaptor sequence was trimmed from raw reads with cutadapt v5.0 (Martin, 2011), and reads were quality filtered with fastp v0.23.4 (Chen, 2023; parameters: -5 20, -3 20, --cut_window_size 5, --cut_mean_quality 15, -q 15) and mapped to the genome with BWA samse (Li, 2013). Variant calling was performed with Freebayes v1.3.6 (Garrison & Marth, 2012; parameters: -m 20, -q 20, -E 3, --min-coverage 20, -V) on merged bam files for each species and data was exported as VCF files. VCFtools (Danecek et al., 2011) was used to reduce variants to bi-allelic SNPs, followed in order by 1) filtering SNPs with a minor allele count of at least 3, 2) dropping individuals that genotyped in fewer than 50% of loci, 3) dropping loci genotyped in fewer than 80% of remaining individuals, and 4) dropping loci with a minor allele frequency of less than 0.01.

Finally, to examine potential for cross amplification in non-Great Lakes *Coregonus* spp., the final subset of panel amplicons were mapped to the European Whitefish *C. lavaretus* sp. ‘Balchen’ genome (GCA_902810595.1, DeKayne et al., 2020), the Amur Whitefish *C. ussuriensis* genome (GCA_031479575.1, Huang et al., 2024), and the Lake Whitefish *C. clupeaformis* genome (GCA_018398675.1, Mérot et al., 2023) with BWA bwasw (Li, 2013). Mapping success of European and Amur whitefishes was compared to mapping success in Lake Whitefish which could be benchmarked with genotyping rates.

### Collection, processing, and GT-seq library preparation of larval and age-1 coregonines in Lake Superior

Collection of larval and age-1 ciscoes in 2019, and age-1 ciscoes in 2020 during annual surveys of Lake Superior was conducted by the U.S. Geological Survey, Great Lakes Science Center (Vinson et al., 2021). Larvae were collected with a paired 1 m^2^, 500 µm mesh neuston net (SEA-GEAR Corporation). The bottom of the net frame was fished ∼0.5 m below the lake surface. The net was fished for 10 minutes at ∼4.0 km per h for ∼0.7 km as determined from the global positioning system on the research vessel. Age-1 ciscoes were collected by bottom trawling using a 12 m-wide Yankee bottom trawl (Vinson et al. 2021). Total lengths (TL, mm) of 2019 larvae were measured in the laboratory. Larvae that could fit within the microscope field of view at the broadest magnification (8x) for the Leica EZ4W were photographed, and TL was measured to the nearest 0.01 mm with calibrated distance lines in Leica Application Suite v3.4.0 (Leica Microsystems; Switzerland). Larvae whose size exceeded the field of view were measured with digital calipers. TL of 2021 larvae was measured to the nearest mm using a standard ruler under a dissecting microscope. TL measurements for 2019-2021 age-1 coregonines were made at the time of collection using a standard ruler at the time of capture.

Preliminary funding was limited for processing 2019 samples, so tissue from up to 12 randomly selected 2019 larval coregonines from each neuston net tow (n=912) was subsampled for GT-seq genotyping. All collected 2019-2021 age-1 coregonines (n=360), and all collected 2021 larval coregonines (n=1,872) were processed. DNA from larval and age-1 samples was extracted using either a 10% Chelex 100 slurry or Qiagen® DNeasy Blood & Tissue kits as described for reference and outgroup coregonines above. Libraries for larval and age-1 samples were prepared following the same protocol described above for panel optimization (Supplemental File S1). The libraries for 2019 larval and age-1 samples were PE 150 sequenced with v1 chemistry on an Illumina NovaSeq platform at the UW Biotechnology Center DNA Sequencing Core Facility (Madison, Wisconsin, USA). The libraries for 2020 and 2021 samples were PE150 sequenced with v2 chemistry on an Illumina MiSeq at the US Geological Survey’s Great Lakes Science Center (Ann Arbor, Michigan, USA).

### GT-seq genotyping and genetic identification of early life coregonines in Lake Superior

Raw sequencing data for references and larval and age-1 coregonines were processed and genotyped in GTscore. Samples with outlying heterozygosity and/or a contamination score>0.50 were dropped before converting genotypes to genepop format. Genotypes were imported into *dartR* (Gruber et al., 2018; Mijangos et al., 2022). First, to determine if Lake, Round, or Pygmy Whitefishes were present in our samples, we reduced the dataset to the final whitefish whitelist and kept individuals genotyped in at least 25% of the loci. We exported reference and unknown data in genepop format and used *assignPOP* (Chen et al., 2018) to identify whitefishes. *C. artedi*, *C. hoyi* and *C. kiyi* references were combined into a single reference population for this analysis. The *assign.X* function was run using the default criterion for retaining the number of principal components (“Kaiser-Guttman”) and the support vector machine (svm) predictive model. Whitefishes with an assignment probability >70% to a reference group were assigned to that group.

Identified whitefishes were removed from the full dataset, and iterative filtering was repeated in *dartR* to generate a dataset for genetic assignment of ciscoes. Filtering steps included removing loci missing in more than 25% of individuals, followed by removing individuals missing more than 62.5% of genotypes. Finally, loci genotyped in less than 75% of the remaining individuals were dropped to generate the final dataset. Genetic similarity of genotyped individuals was examined with principal components analysis (PCA) using the glPCA function before performing genetic assignments.

Preliminary genetic assignments were performed with *assignPOP* (Chen et al., 2018) following the same methods as described for whitefish identification. Final genetic assignments were generated with model-based clustering in STRUCTURE v2.3.4 (Pritchard et al., 2000; Falush et al., 2003, 2007). Given the large number of assignments to be performed, our dataset was randomly subsampled to batches of ∼100 unknowns to be run with the reference individuals (n=129). STRUCTURE was run without prior population information and with the admixture model and correlated allele frequencies using a burn-in of 50000 followed by 100000 MCMC steps. Cluster analysis on each batch was run for K=3 with 5 replicates. Replicates were merged with the R package *pophelper* (Francis 2017), and larvae and juveniles with Q-scores >70% to a reference group cluster were assigned to that cluster following Ackiss et al. (2020). Individuals that did not assign with high probability to a single cluster were labeled as putative hybrids.

### Preliminary analysis of early life coregonines in Lake Superior

Identified larvae from 2019 and 2021 were plotted to show the composition of species at each collection location in Lake Superior. The spatial (whole lake) and temporal (May 29 - July 23) spread of larval collections was broadest in 2019 because the extent of these surveys was not impacted by the restrictions introduced during the COVID pandemic in 2021 collections. Therefore, we used the 2019 cohort data to examine changes in total length (TL) over time. We extrapolated larval growth rate per day (mm/day) using linear regression for the first month of collections which are more likely to represent a demographic cohort. Identified age-1s from 2019-2021 were also plotted to show the composition of species at each collection location, and genetic identifications were used to assess the accuracy of field identifications in 2019-2020 age-1s.

## Results

### Development of an amplicon panel from RAD-seq data

A total of 858 samples from the three major extant species in the *Coregonus artedi* species complex representing all the lakes with existing populations (Figure 1) were genotyped at 49,841 SNPs using RAD-seq. Ten initial reporting groups were identified with a threshold of *G*_ST_ ≥0.15 (Supplemental File 1, Supp. Fig.1A). Results indicated a large proportion of individuals from Grand Portage in Lake Superior (A_SGP, 33%) and the North Channel and Georgian Bay regions in Lake Huron (A_HNC, 25%) assigned to nearby clusters – potentially because these collections were comprised of non-spawning period fish. These locations were combined with their counterparts to generate eight final reporting groups (Supplemental File 1, Supp. Fig.1B; Table 1), including one for each species in Lake Superior, one for *C. hoyi* from Lake Michigan and Lake Huron, one for Lake Michigan *C. artedi*, two for Lake Huron *C. artedi*, and one for Lake Ontario *C. artedi* (Fig. 1). The results from candidate panels indicated similar performance between a panel comprised of 500 high *G′*_ST_ SNPs among Lake Superior ciscoes and 100 SNPs with high basin-wide *H*_e_ and a panel comprised of both 500 high *G′*_ST_ SNPs and 100 SNPs with high *H*_e_ from Lake Superior ciscoes (Supplemental File 1, Supp. Table 1, Supp. Fig. 2). We selected the latter for designing primers along with the 45 high *F*_ST_ transcriptomic loci.

The original 645 GT-seq panel loci were iteratively reduced over three rounds of test amplification. Over-represented loci were identified and removed after each round of test amplification resulting in a final panel of 494 loci composed of 467 RAD-seq based loci and 27 RNA-seq based loci containing 1220 target SNPs. A total of 434 panel loci (87.5%) mapped with 2 or fewer mismatches to the *C. artedi* genome across all 38 chromosomes (Fig. 2).

**Figure 2.**
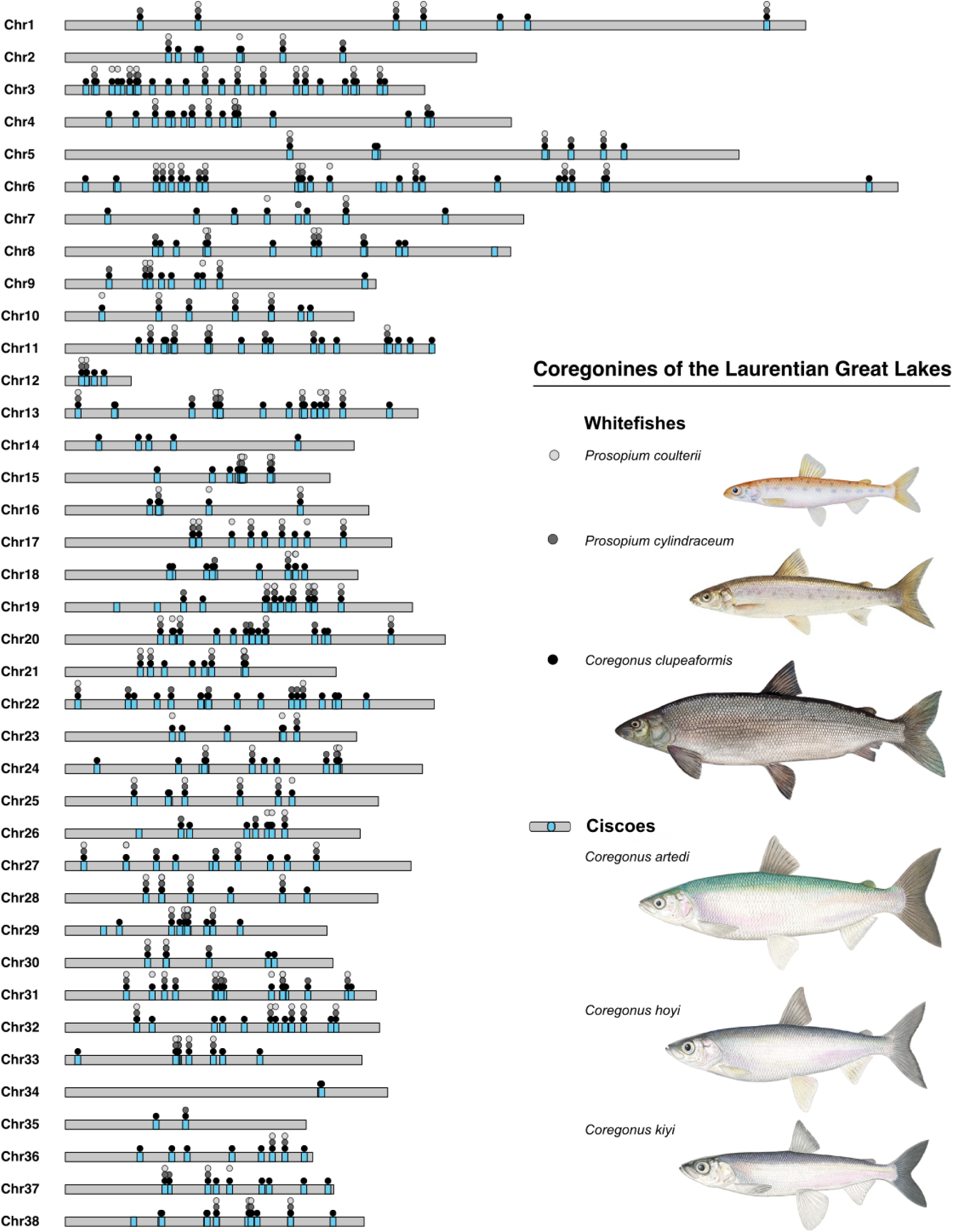
Cross amplification in Great Lakes coregonines and location of the GT-seq panel loci in the *C. artedi* genome. Loci that cross-amplified in whitefishes are annotated with circles – Pygmy Whitefish *P. coulterii* (light gray), Round Whitefish *P. cylindraceum* (dark grey), Lake Whitefish *C. clupeaformis* (black).

### Quantifying cross-amplification success, and modifying probes for outgroup assignment

After reviewing cross-amplifying loci, existing panel probes were modified to account for 22 additional whitefish variants bringing the total number of target SNPs to 1242 (Supplemental File 1, Supp. Fig. 3; Supplemental Files 3-4). Lake Whitefish *C. clupeaformis* genotyped at 464 panel loci (93.9%) at 1101 of the target SNPs with an average genotype rate of 93.9% (range 0.51-100%). Round Whitefish *P. cylindraceum* genotyped at 205 (41.5%) of panel loci at 404 of the target SNPs with an average genotype rate at these loci of 85.3% (range 0.50-97.4%). Pygmy Whitefish *P. coulterii* genotyped at 186 (37.7%) of panel loci at 338 of the target SNPs with an average genotype rate at these loci of 75.9% (range 0.50-90.6%). Cross-amplifying loci in Lake Whitefish mapped to all 38 chromosomes of the *C. artedi* genome. Cross-amplifying loci in Round and Pygmy Whitefish mapped to 36 and 35 chromosomes, respectively. Locations of cross-amplifying loci in whitefishes are annotated in Fig. 2.

A whitelist of 571 polymorphic SNPs across all whitefishes and a whitelist of 428 polymorphic SNPs in *Prosopium* whitefishes performed with 100% assignment accuracy for Lake Whitefish, Round Whitefish, and Pygmy Whitefish at all combinations of training data and loci. Since *Prosopium* whitefishes generally cross-amplify in fewer loci, we proceeded with the 428 SNP *Prosopium* whitefish-specific whitelist (Supplemental File 5) in downstream analyses to reduce the potential for losing samples through the missing data (>25% loci) filter. The assignment accuracy plot and a PCA of reference individuals using the 428 SNP whitelist is reported in Supplement File 1 (Supp. Figs. 4, 5).

When sequenced amplicons were mapped to the *C. artedi* genome and variants were called and filtered within species, Lake Whitefish produced a dataset of 1280 bi-allelic SNPs with a genotype rate of >80% across 56 individuals. Round Whitefish and Pygmy Whitefish produced datasets of 124 and 297 bi-allelic SNPs with a genotype rate of >80% across 34 and 22 individuals, respectively. A summary of GT-seq panel cross-amplification in related Great Lakes coregonines is presented in Table 2.

**Table 2.** Cross-amplification of the GT-seq panel on other Great Lakes coregonines. Summary includes the number (percent) of the 494 panel loci that cross-amplified, the number (percent) of the genotyped SNPs called from GT-seq panel probes, the number of chromosomes of *C. artedi* genome cross-amplifying loci were found in (number of chromosomes to which panel amplicons mapped in the species’ respective genome), and the number of biallelic SNPs within species after filtering when amplicons were mapped to the *C. artedi* genome with the number (n) of genotyped individuals.

| Species | Common Name | GT-seq Panel |  |  | Genome mapped |
| --- | --- | --- | --- | --- | --- |
|  |  | Amplified Loci | Genotyped SNPs | Chrom | Filtered SNPs (n) |
| *C. clupeaformis* | Lake Whitefish | 464 (93.9%) | 1101 (90.2%) | 38 (39) | 1280 (56) |
| *P. cylindraceum* | Round Whitefish | 205 (41.5%) | 404 (33.1%) | 36 (-) | 124 (34) |
| *P. coulterii* | Pygmy Whitefish | 186 (37.7%) | 338 (27.7%) | 35 (-) | 297 (22) |

Finally, mapping panel amplicons to whitefish genomes suggests cross-amplifying loci are widespread across the genome. When amplicons were mapped to the European Whitefish *C. lavaretus* sp. ‘Balchen’ genome, 411 panel loci (83.2%) had a mapping quality (MAPQ) score of ≥30 to regions in 39 out of 40 chromosomes. No matches occurred in chromosome 40, the smallest chromosome in the *C. lavaretus* genome at ∼1 Mbp. When mapped to the Amur Whitefish *C. ussuriensis* genome, 392 panel loci (79.4%) had a MAPQ score ≥30 to regions in 39 out of 40 chromosomes and one scaffold. Similarly to European Whitefish, no matches occurred in chromosome 40, which is also ∼1 Mbp in *C. ussuriensis.* Comparatively, while 464 panel loci (93.9%) successfully amplified in Great Lakes Lake Whitefish *C. clupeaformis* samples, only 404 loci (81.8%) mapped with a MAPQ score of ≥ 30 across 39 of 40 chromosomes (no matches in chromosome 38 of the *C. clupeaformis* genome, ∼11 Mbp) and 10 scaffolds suggesting mapping cisco-based amplicons could underestimate cross amplification potential.

### GT-seq genotyping and genetic identification of early life coregonines in Lake Superior

Adult references used for assignments exhibited strong clustering by species. For example, the average Q-scores for references to their associated species cluster in the first of 34 subsetted STRUCTURE runs were *C. artedi* = 0.90, *C. hoyi* = 0.92, and *C. kiyi* = 0.95 (STRUCTURE plot in Supplemental File 1, Supp. Fig. 6). Clear differentiation between reference ciscoes can also be seen when the dataset is reduced to the whitefish whitelist (Supplemental File 1, Supp. Fig. 5). For all collections of larvae and age-1s, more than 90% of samples were able to be identified with the GT-seq panel (91.7-100%, Table 3). Of the larvae collected in 2019, most were *C. artedi* (n=569), followed by *C. kiyi* (n=179) and *C. hoyi* (n=5). Genetic species identifications of larvae collected in 2021 showed a similar proportion (*C. artedi*, n=1522; *C. kiyi*, n=130; *C. hoyi*, n=16). Putative hybrids, those individuals which generated a Q-score below our 0.70 threshold to assign to a cluster, comprised 11.5% and 9.7% of genotyped larvae in 2019 and 2021, respectively. In the age-1 collections from 2019-2021, most were *C. hoyi* (n=235) followed by a similar number of *C. artedi* (n=39) and *C. kiyi* (n=31). Putative hybrids comprised 4.2-16.0% of genotyped age-1s. Additionally, the whitefish whitelist identified 13 Lake Whitefish in the larval collections and 2 Pygmy Whitefish in the 2020 age-1 collection. A summary of identified larvae and age-1s is presented in Table 3, *assignPOP* assignment scores and STRUCTURE Q-scores are reported in Supplemental File 6.

**Table 3.** Genetic identifications of early life coregonines collected in Lake Superior 2019-2021. Proportion of putative hybrids (i.e. fishes whose scores were below our assignment threshold) in each collection is listed as a percentage of genotyped individuals.

| Species | 2019 |  | 2020 | 2021 |  |
| --- | --- | --- | --- | --- | --- |
|  | larvae | age-1s | age-1s | larvae | age-1s |
| *C. artedi* | 569 | 4 | 0 | 1522 | 35 |
| *C. kiyi* | 179 | 17 | 3 | 130 | 11 |
| *C. hoyi* | 5 | 17 | 64 | 16 | 154 |
| *C. clupeaformis* | 10 | 0 | 0 | 3 | 0 |
| *P. coulterii* | 0 | 0 | 2 | 0 | 0 |
| putative hybrid | 99 (11.4%) | 6 (13.6%) | 3 (4.2%) | 179 (9.7%) | 38 (16.0%) |

### Preliminary analysis of early life coregonines in Lake Superior

Investigation of the spatial distribution of identified coregonines indicated that *C. artedi* and *C. kiyi* are broadly distributed in the surface waters throughout Lake Superior in spring and early summer (Fig. 3). The small number of *C. hoyi* larvae collected were in the nearshore areas of the western arm of the lake. The 10 Lake Whitefish larvae caught in 2019 were collected mid-June to mid-July nearshore along the northern coastline of Lake Superior, and the three Lake Whitefish larvae caught in 2021 were collected May 20 and 21 nearshore in the western arm of the lake. Only five *C. hoyi* larvae were collected in 2019, so broad comparisons of larval size by collection date were limited to *C. artedi* and *C. kiyi*.

**Figure 3.**
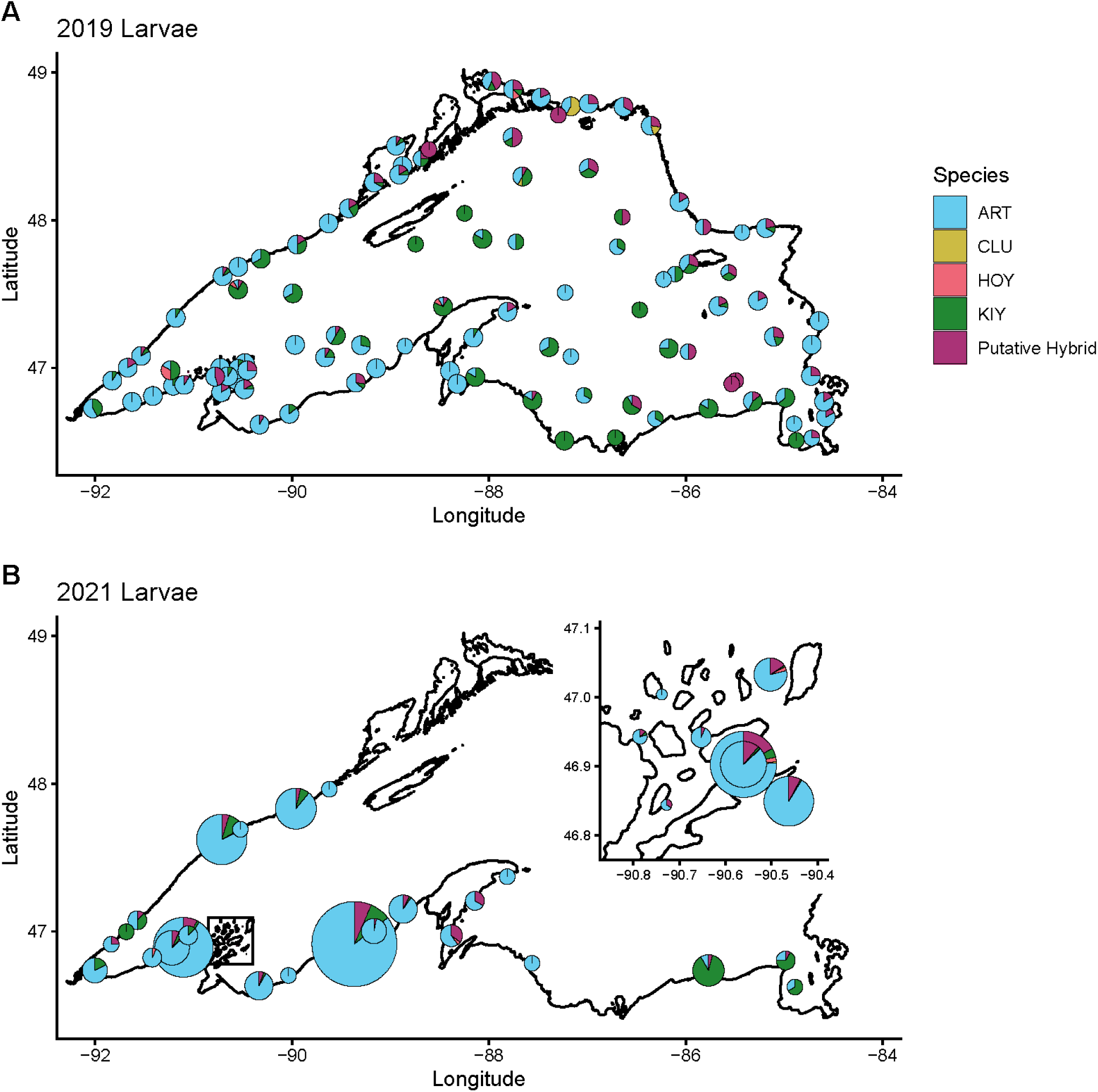
Spatial distribution of 2,712 genetically identified larvae collected from Lake Superior in 2019 (A) and 2021 (B). The extent of 2021 larval collections was limited due to travel restrictions during the COVID pandemic. Collections from this year occurred largely in U.S. waters in the region near the Apostle Islands (inset, panel B). Pie chart size is proximal to number of larvae from each collection site. Abbreviations: ART – *C. artedi* (Cisco), CLU - *C. clupeaformis* (Lake Whitefish), HOY – *C. hoyi* (Bloater), and KIY – *C. kiyi* (Kiyi).

The median total length (TL) of *C. artedi* and *C. kiyi* larvae in 2019 was 12.97 mm (range: 8.87-21.04 mm) and 14.19 mm (range: 10.19-17.42 mm), respectively. Of the smaller catches, the median TL of the five *C. hoyi* larvae was 14.08 mm (range: 12.42-14.88 mm), and the median TL of the 10 Lake Whitefish larvae was 16.04 mm (range: 13.42-21.23 mm). Regression analysis indicated an estimated growth rate of 0.11mm/day for *C. artedi* and 0.06mm/day for *C. kiyi* from the period of late May to late-June 2019 (Fig. 4). The median TL of *C. artedi* and *C. kiyi* larvae in 2021 was 12 mm (range: 8-20 mm) and 13 mm (range: 11-20 mm), respectively. Of the smaller catches in 2021, the median TL of the 16 *C. hoyi* larvae was 10 mm (range: 9-12 mm), and the median TL of the three Lake Whitefish larvae was 16.04 mm (range: 13.42-21.23).

**Figure 4.**
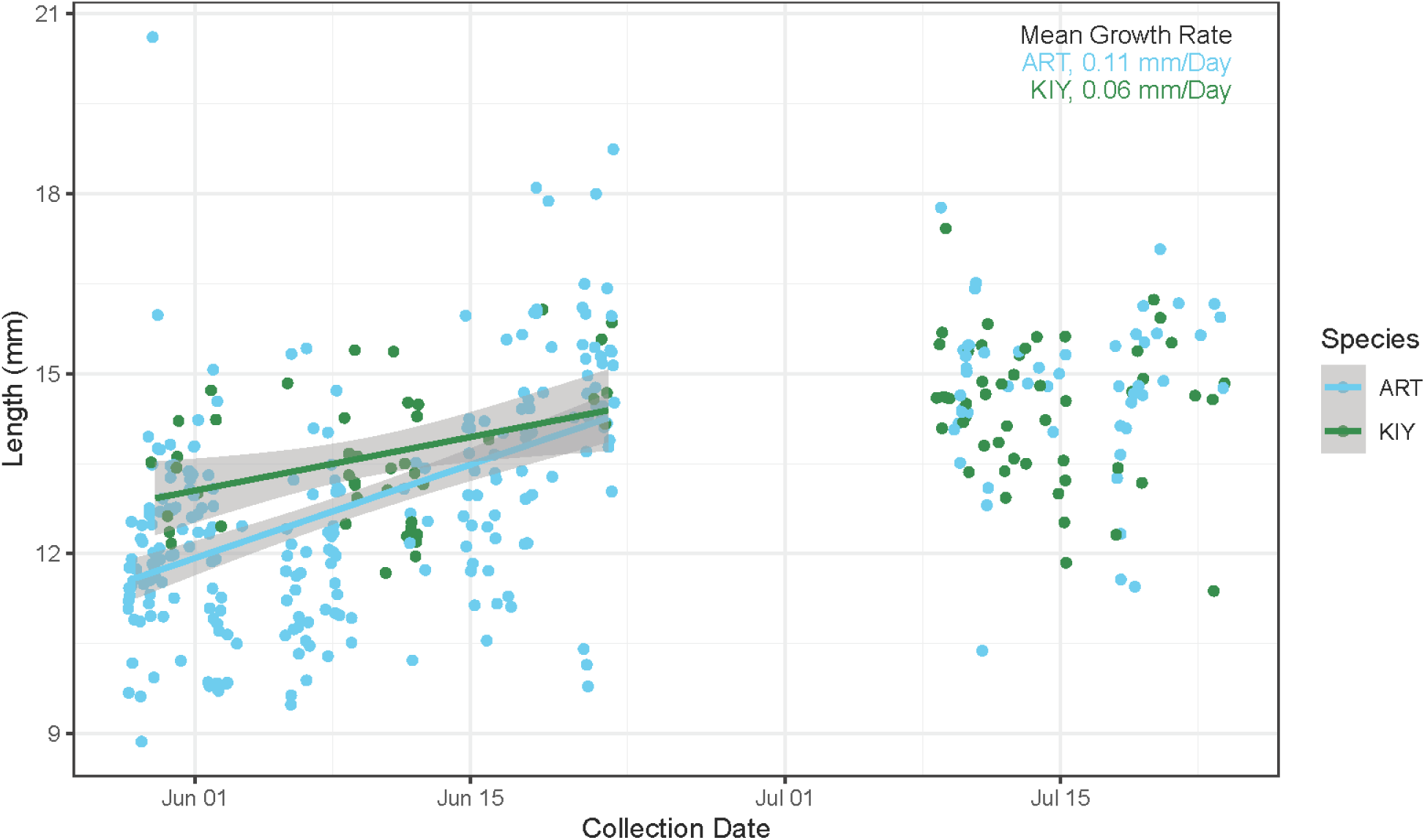
Estimated growth rates of 569 *C. artedi* (ART) and 179 *C. kiyi* (KIY) larvae from late May-July 2019.

Comparisons of field and genetic identifications of age-1s highlight the challenges of identifying early life coregonines (Fig. 5). Of the three major species, *C. kiyi* was most likely to be correctly identified in the field. All age-1s identified as *C. kiyi* genotyped as *C. kiyi*. The 2 Pygmy Whitefishes collected with the 2020 age-1s were caught in the Apostle Islands and originally identified as “unknown coregonines” as their total length (25 mm, 26 mm) was significantly smaller than the average for age-1 ciscoes (in 2020, 137.6 mm). Evaluating age-1 species identification by collection location indicated that the distribution of juvenile *C. kiyi* is largely offshore and the distribution of juvenile *C. artedi* and *C. hoyi* is more nearshore (Supplemental File 1, Supp. Fig. 7).

**Figure 5.**
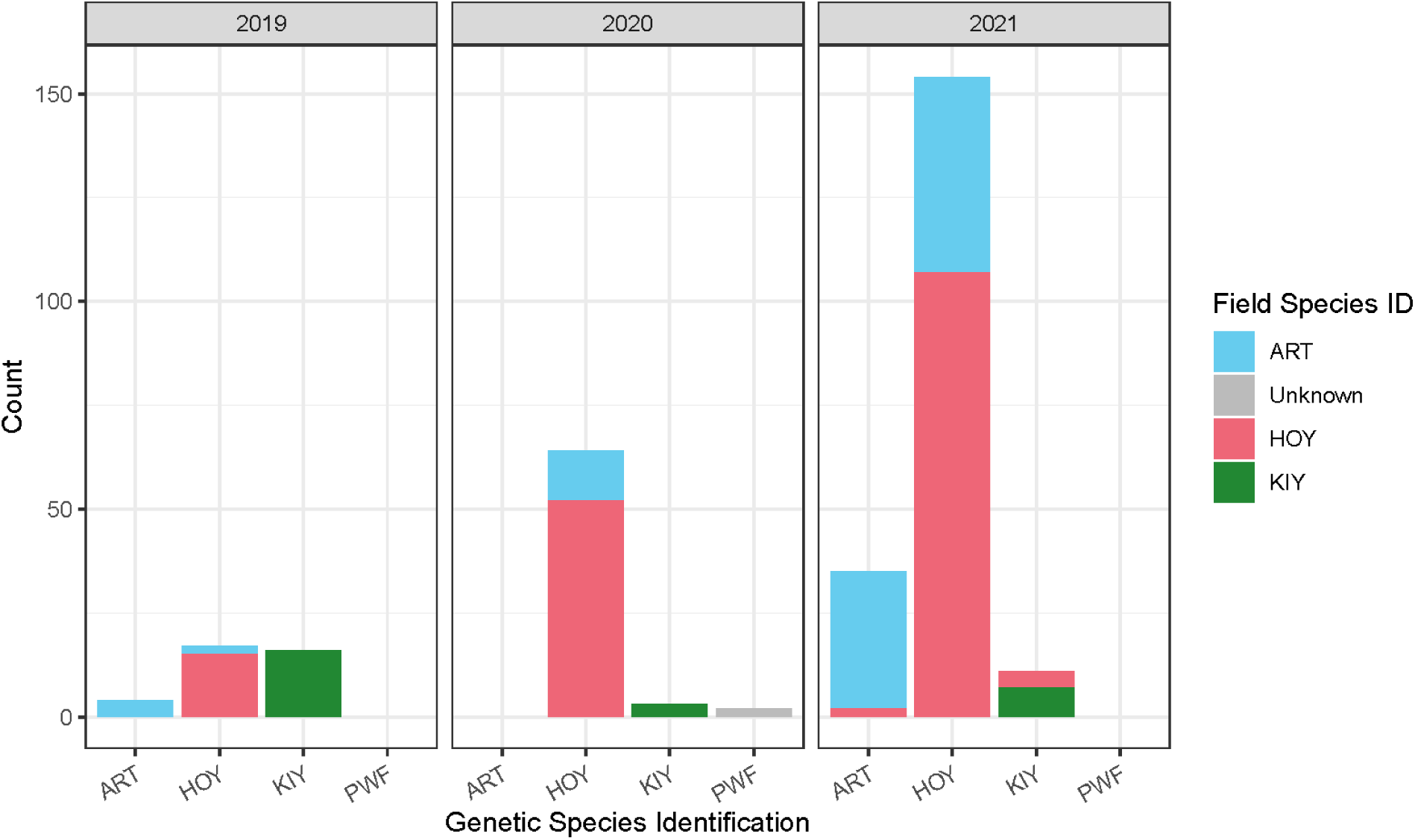
Comparisons of field versus genetic species identifications from 2019-2021 age-1 collections. Abbreviations: ART – *C. artedi* (Cisco), HOY – *C. hoyi* (Bloater), KIY – *C. kiyi* (Kiyi), PWF – *P. coulterii* (Pygmy Whitefish).

## Discussion

### Development of a GT-seq panel for cryptic species identification

We built a flexible GT-seq panel of 494 loci containing 1242 target SNPs from RAD- and RNA-seq data capable of identifying distinct *Coregonus* species that arose through an evolutionarily recent radiation in the Laurentian Great Lakes as well as congener and confamilial coregonines. A cost-effective GT-seq panel opens the door to a myriad of potential research that can investigate species-specific early life history and represents a significant improvement over the two methods used to genetically identify early life coregonines in the Great Lakes prior to the development of this GT-seq panel. In Lake Ontario, the cytochrome oxidase subunit I (COI) gene was used to identify 726 Cisco *Coregonus artedi* and Lake Whitefish *C. clupeaformis* larvae collected in 2014 (George et al., 2018). While variation in COI and other mitochondrial regions can distinguish between ciscoes and whitefishes, it cannot distinguish ciscoes from each other (Bernatchez et al., 1991; Turgeon & Bernatchez, 2003; Lachance et al., 2021) which is a common challenge for evolutionarily young species (van Velzen et al., 2012). This method was a viable option in Lake Ontario at the time because a century of anthropogenic impacts had reduced the diversity of ciscoes from four species to one (Eshenroder et al., 2016). However, a bi-national restoration initiative has been underway since 2012 to reintroduce Bloater *C. hoyi* to Lake Ontario (OMNRF 2015; Weidel et al., 2022), and as this effort progresses, COI may no longer be an accurate method for early life stage identification. Additionally, a 2018 study examining emergence and species composition of larval coregonines in the Apostle Islands of Lake Superior used COI to differentiate Lake Whitefish from ciscoes, then genetically identified 193 larvae that barcoded as ciscoes with RAD-seq (Lachance et al., 2021). Reduced representation sequencing methods such as RAD-seq can be extremely effective at identifying cryptic diversity (Pante et al., 2015; Herrera & Shank, 2016) but can cost up to 5x more per sample (Meek & Larson, 2019) significantly limiting the scope of study. Here, we demonstrate the relative ease with which thousands of larvae can be identified with a single, cost-effective method, enabling research and monitoring that may rely on broad or repeated sampling to achieve robust results.

### Cross amplification across coregonine species

While originally designed to distinguish members of the cisco species complex, we were able to adapt SNP probes for genetic identification of Great Lakes whitefishes with relative ease and high accuracy. Cross amplification was more successful between species within genus than between genera within family echoing broad patterns observed in microsatellite cross amplification in fishes and other large taxonomic groups (Barbará et al., 2007). Despite the reduction in success across genera, cross amplification was widespread across the Cisco *C. artedi* genome in both congeneric and confamilial coregonines. Genomic resources for coregonines are still emergent - only two North American species (Cisco *C. artedi* and Lake Whitefish *C. clupeaformis*) have published genomes (Backenstose et al., 2024; Merot et al., 2023). Synteny between their chromosomes has not been evaluated in the published literature; however, the high proportion of loci that cross-amplify in Lake Whitefish across the 38 resolved chromosomes of the *C. artedi* genome and good quality amplicon mapping in 39 or 40 Lake Whitefish chromosomes indicates these loci remain widespread across the *C. clupeaformis* genome. High synteny is present between the Lake Whitefish and the European Whitefish *C. lavaretus* sp. Balchen genomes (Mérot et al., 2023) and similarly between *C. lavaretus* sp. Balchen and the Amur Whitefish *C. ussuriensis* in Asia (Huang et al., 2024). Relatively comparable proportions of panel amplicons mapping with good quality to the genomes of these species and Lake Whitefish indicates synteny could be used as a preliminary predictor for cross amplification success. The only chromosome that panel amplicons did not map to in European and Amur Whitefish is a small, ∼ 1 Mbp ‘super-scaffold’ (Chromosome 40; De-Kayne et al., 2020, Huang et al., 2024) detected in both species that De-Kayne et al. (2020) hypothesized could be part of another chromosome. Our results suggest that panel loci may remain broadly representative of neutral or adaptive genomic variation in these species and could be successfully mined for off target use in other *Coregonus* spp. across the northern hemisphere.

Cross amplification was lower in the *Prosopium* whitefishes, though the smaller number of loci that amplified in these fishes remained widespread across the *C. artedi* genome. Despite the decrease in loci, we were still able to achieve high accuracy in genetic species identification for both Round Whitefish *P. cylindraceum* and Pygmy Whitefish *P. coulterii* from the Great Lakes. No genomes have been published for any *Prosopium* spp. so inferences from synteny cannot be made as to whether cross-amplifying loci may be widespread across independent *Prosopium* chromosomes. Karyotypes for *P. cylindraceum* and *P. coulterii* indicate nearly the same number of chromosomes (2n = 78 and 82, respectively; Booke 1968) as in ciscoes (2n=80; Booke, 1968; Phillips et al., 1996); however, *P. cylindraceum* and *P. coulterii* are the most ancestral of the genus (Vuorinen et al., 1998), and karyotypes of the four other *Prosopium* spp. show successively reduced numbers of chromosomes (2n=64-74; Booke, 1970, 1974) from Robertsonian translocations facilitating the process of rediploidization and speciation (Booke, 1974; Phillips & Rab, 2001). Whether the approximately 200 panel loci that cross-amplified in *P. cylindraceum* and *P. coulterii* would continue to successfully amplify across the breadth of *Prosopium* spp. in the face of significant chromosomal rearrangements remains to be tested.

### Early life coregonines in Lake Superior

The distributions of early life coregonines appear heavily influenced by mobility and emergence timing. In Lake Superior, pelagic *C. artedi* spawn in nearshore waters of depths between 15-30 m (Koelz, 1929; Yule et al., 2006), while deepwater *C. kiyi* spawn offshore in depths >100m (Vinson et al., 2023). Larvae of both species were consistently collected in the upper meter of the lake in both near and offshore tows, suggesting larval orientation is largely driven by yolk-sac buoyancy, positive phototaxis, and passive dispersal via currents. Conversely, age-1s of these species segregated to nearshore midwater (*C. artedi*) and offshore deepwater (*C. kiyi*) following the known species-specific distributions of juveniles and adults of these species (Stockwell et al., 2006; Yule et al., 2013; Rosinski et al., 2020). The fact that *C. hoyi* was rarely collected in larval tows from May-July likely reflects spawn and emergence timing. The three species spawn in consecutive months in the fall and winter, beginning with *C. artedi* (November, Koelz, 1929), followed by *C. kiyi* (late December-late January; Vinson et al., 2023), and then *C. hoyi* (February-March, Dryer & Beil, 1968). Continued sampling into August and September or at greater water column depths could better characterize the distribution of newly emerged *C. hoyi* larvae to confirm if their distributional patterns mimic that seen in *C. artedi* and *C. kiyi*.

Growth rates and larval size indicated species-specific differences between *C. artedi* and *C. kiyi* larvae. Notably, *C. artedi* larvae were consistently caught at smaller sizes than *C. kiyi* larvae in both 2019 and 2021. Whether this is reflective of a smaller size at hatch for *C. artedi*, growth that may occur between hatch in deeper water and presence in surface waters of *C. kiyi*, or some other physiological difference is unknown. Hatchery collections of *C. artedi* have been established for stocking purposes since the late 1800s (McKenna et al., 2025) and could be used to more comprehensively examine larval development. However, the rearing of *C. kiyi* eggs has only been documented once (Todd et al., 1981) as the winter spawning period and depth of *C. kiyi* makes collection of spawning individuals notoriously difficult. Vinson et al. (2023) were able to collect spawning *C. kiyi* from Lake Superior in 2017-2019 and determined that average egg diameter in *C. kiyi* (∼1.7 mm) was not significantly different than *C. artedi* (Koenigbauer et al., 2022). Despite the presence of smaller *C. artedi* larvae in surface collections, higher growth rates were observed in *C. artedi* than in *C. kiyi*, though the rates presented here should be viewed cautiously. Adult *C. artedi* is substantially larger than adult *C. kiyi* (Koelz, 1929) so a faster growth rate would not be unexpected; however, more robust estimates than those presented here could be obtained in the future by better controlling for geographic region across time and potential gear bias on larval size.

The presence of putative hybrids in larval and age-1s samples reflects evidence for hybridization previously seen in coregonines across the northern hemisphere. Hybridization and speciation reversal driven by anthropogenic impacts like eutrophication and non-native species introduction has been documented among *Coregonus* spp. in number of European Alpine and subarctic lakes (Kahilainen et al., 2011; Vonlanthen et al., 2012; Frei et al., 2023). Hybridization has also been seen between species pairs of Lake Whitefish *C. clupeaformis* in Canadian lakes and has been hypothesized to be associated with incipient ecological speciation (Lu et al., 2001; Rougeux et al., 2017). Recently with both RAD-seq and whole-genome sequencing data, hybrids were documented in Lake Superior (Ackiss et al., 2020; Lachance et al., 2021, Backenstose, 2024). Unlike many of the Alpine and subarctic lake populations, the nature of hybridization among ciscoes in Lake Superior is unstudied. The average proportion of putative larval and age-1s hybrids detected in our study (∼11%) was similar to the hybrid rate previously found in adults (∼9%; Ackiss et al., 2020) but was nearly triple the proportion of hybrid larvae found in Lachance et al. (∼4%; 2021). The increase in putative hybridization rates in larvae detected here may be the result of a more than seven-fold increase in sample numbers, a reflection of stochastic differences between year classes, or evidence of ecological change breaking down pre-zygotic isolating mechanisms. Long term monitoring in Lake Superior of both larvae and adults could illuminate whether the proportions of putative hybrids are indicative of natural variation in an evolutionarily recent ecological species radiation that is undergoing speciation-with-gene flow or evidence that ecosystem change may be driving speciation reversal.

Comparisons of field and genetic identifications of age-1s reflected the challenges that cryptic early life stages can present to accuracy in identifications. Misidentifications of age-1 *C. kiyi* were relatively rare, with only a few individuals from 2021 originally identified as *C. hoyi*. Misidentifications of this type are not uncommon in adults as well. Early descriptions of the species complex noted *C. hoyi* and *C. kiyi* could be confused with each other in Lake Superior (Koelz, 1929), and genetic analysis of adults in the Apostle Islands found three out of 24 individuals originally identified as *C. hoyi* genotyped as *C. kiyi* (Ackiss et al., 2020). In contrast, no misidentifications between adult *C. artedi* and *C. hoyi* were seen by Ackiss et al. (2020), and these were the most common misidentifications in age-1s here. This may be an indication of variance between individuals performing identification or reduced accuracy or consistency in the characters used to distinguish the two in early life. With new accessibility to a tool for genetically confirming species identification, characters for morphological identification of age-1s can be reevaluated to increase accuracy.

These preliminary results illustrate the value of a simple, targeted, and cost-effective solution for cryptic early life species identification, particularly when species-specific outcomes can be variable.

While all Great Lakes coregonines have experienced significant population declines, the deepwater species of the *C. artedi* species complex were inordinately impacted by the rapid ecological changes that have occurred over the past century (Eshenroder et al., 2016). More recently, however, declines in shallow water Lake Whitefish *C. clupeaformis* have been observed (Ebener et al., 2021; Cunningham & Dunlop, 2023) while some shallow water *C. artedi* populations appear to be stable or increasing (Tingley et al., 2025). Early life in fishes is a critical period underpinned by rapid changes in morphology, ecology, and habitat use, and high mortality (Chambers & Trippel, 2012). A better understanding of the early life period could play a crucial role in determining the nature of differential species responses to ecosystems undergoing environmental or ecological change. This panel now provides a cost-effective method for future research of the *C. artedi* species complex at the species-specific level across the whole life history spectrum, thus opening the door to new insights into the mechanisms by which species-specific adaptation has driven speciation and outcomes in Great Lakes coregonines.

## Supporting information

Supplemental File 2 GTseq Lib Protocol

Supplemental File 3 Probe File

Supplemental File 4 Primer Sequences

Supplemental File 5 Whitefish Whitelist

Supplemental File 6 Genetic IDs and Q-scores

## Acknowledgements

The authors would like to thank Jared Homola and Nicholas Sard for their reviews of an earlier version of this manuscript. The authors would also like to thank Tommy Hill for photographing, measuring, and extracting DNA from 2019 larval samples and extracting DNA from Lake Michigan Lake Whitefish reference samples, and Sydney Phillips for measuring and pre-processing 2021 larval samples. Samples used in this work were collected by the Great Lakes Fishery Commission, Ontario Ministry of Natural Resources, Chippewa Ottawa Research Authority, Sault Ste. Marie Tribe, Little Traverse Bay Band of Odawa Indians, Wisconsin Department of Natural Resources, United States Fish and Wildlife Service, and the United States Geological Survey. Illustrations of coregonines used in Figure 2 are credited as follows: Pygmy Whitefish (Paul Vecsei), Lake Whitefish (Cory Brant), Round Whitefish (Julia Ellen Edmonson), and Ciscoes (Paul Vecsei, reproduced from Eshenroder et al. 2016 with permission). Any use of trade, firm, or product names is for descriptive purposes only and does not imply endorsement by the U.S. Government.

## Authors’ Contributions

**ASA** – RAD-seq library preparation and genotyping, GT-seq panel development/optimization, GT-seq genotyping, data analysis, lead manuscript development/writing; **MRV** – project conception, collected samples, manuscript writing/editing; **AJR** – performed sequencing, data analysis, data archiving, manuscript writing/editing; **KMG** – supported GT-seq panel development/optimization, manuscript editing; **TJK** - provided transcriptomic data, manuscript writing/editing; **JVS** – performed DNA extraction, GT-seq library preparation, data analysis; **WS** – project conception, collected samples, manuscript editing; **DLY** – collected samples, manuscript writing/editing; **WAL** – project conception, manuscript editing. All authors approved the final draft submitted for publication.

## Data Accessibility

RAD sequence data from *C. artedi*, *C. hoyi*, and *C. kiyi* from the Apostle Islands and *C. artedi* from Boot Island were previously archived on NBCI sequence read archive (Project IDs: PRJNA600818, PRJNA1266121). Sample and sequence data for all samples genotyped with RAD-seq and GT-seq can be found on ScienceBase (https://doi.org/10.5066/P13W68YB, https://doi.org/10.5066/P145DC3U). RNA-seq reads from *C. artedi*, *C. hoyi*, and *C. kiyi* from Lake Superior were previously archived on NCBI sequence read archive (Project ID PRJNA659559).

