## Supplemental File 1 Tables Figures for "A genomic tool to tackle cryptic diversity demonstrates the potential for off-target use of GT-seq panels"

**A genomic tool to tackle cryptic diversity demonstrates the potential for off-target use of GT-seq panels**

Amanda S. Ackiss, Mark R. Vinson, Ann J. Ropp, Kristen M. Gruenthal, Trevor J. Krabbenhoft, Joseph V. Siegel, Wendylee Stott, Daniel L. Yule, Wesley A. Larson

Disclaimer: Any use of trade, firm, or product names is for descriptive purposes only and does not imply endorsement by the U.S. Government.

**Supplemental Table 1.** The eight RADseq locus combinations used for simulating panel performance. Combinations targeted the top loci with the highest differentiation (*G′*_ST_) or diversity (*H*_e_) estimates with both SNP and haplotype datasets. Differentiation and diversity estimates were generated for two datasets: 1) Lake Superior individuals, representing the three major extant species, and 2) all individuals (basin-wide). The locus combination in **bold** was chosen for panel development.

| **Total # loci** | **Locus combination** | **Variant type** | **Assignment Accuracy** |
| --- | --- | --- | --- |
| 600 | 500 *G′*_ST_ (Superior), 100 *H*_e_ (basin-wide) | SNP | 0.977 |
| **600** | **500 *G****′***_ST_ (Superior), 100 *H*_e_ (Superior)** | **SNP** | **0.977** |
| 600 | 600 *G′*_ST_ (basin-wide) | Haplotype | 0.969 |
| 600 | 600 *G′*_ST_ (Superior) | Haplotype | 0.970 |
| 600 | 600 *H*_e_ (basin-wide) | Haplotype | 0.958 |
| 600 | 600 *H*_e_ (Superior) | Haplotype | 0.957 |
| 600 | 600 *G′*_ST_ (Superior) | SNP | 0.971 |
| 600 | 600 *G′*_ST_ (basin-wide) | SNP | 0.963 |

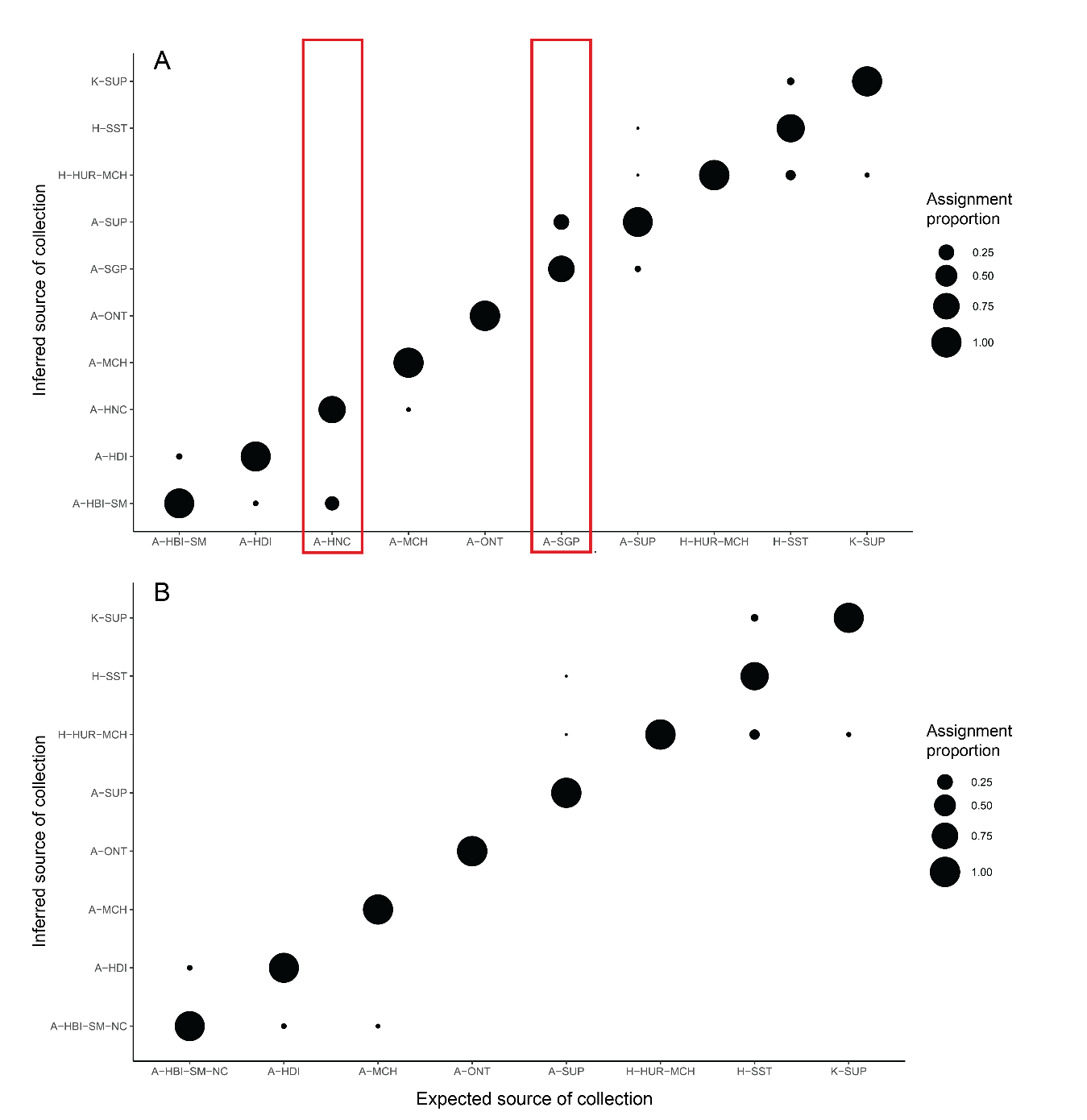

**Supplemental Figure 1**. Self-assignment proportions of reporting groups. Panel A: Preliminary assignment tests indicated that two groups – Grand Portage (A-SGP) and the North Channel/Georgian Bay region (A-HNC) – had more than 25% of individuals assign to other locations (red bars). Panel B: Final reporting groups used to assess candidate panels, with individuals from A-SGP and A-HNC combined with closely associated groups (A-SUP and A-HBI-SM-NC, respectively).

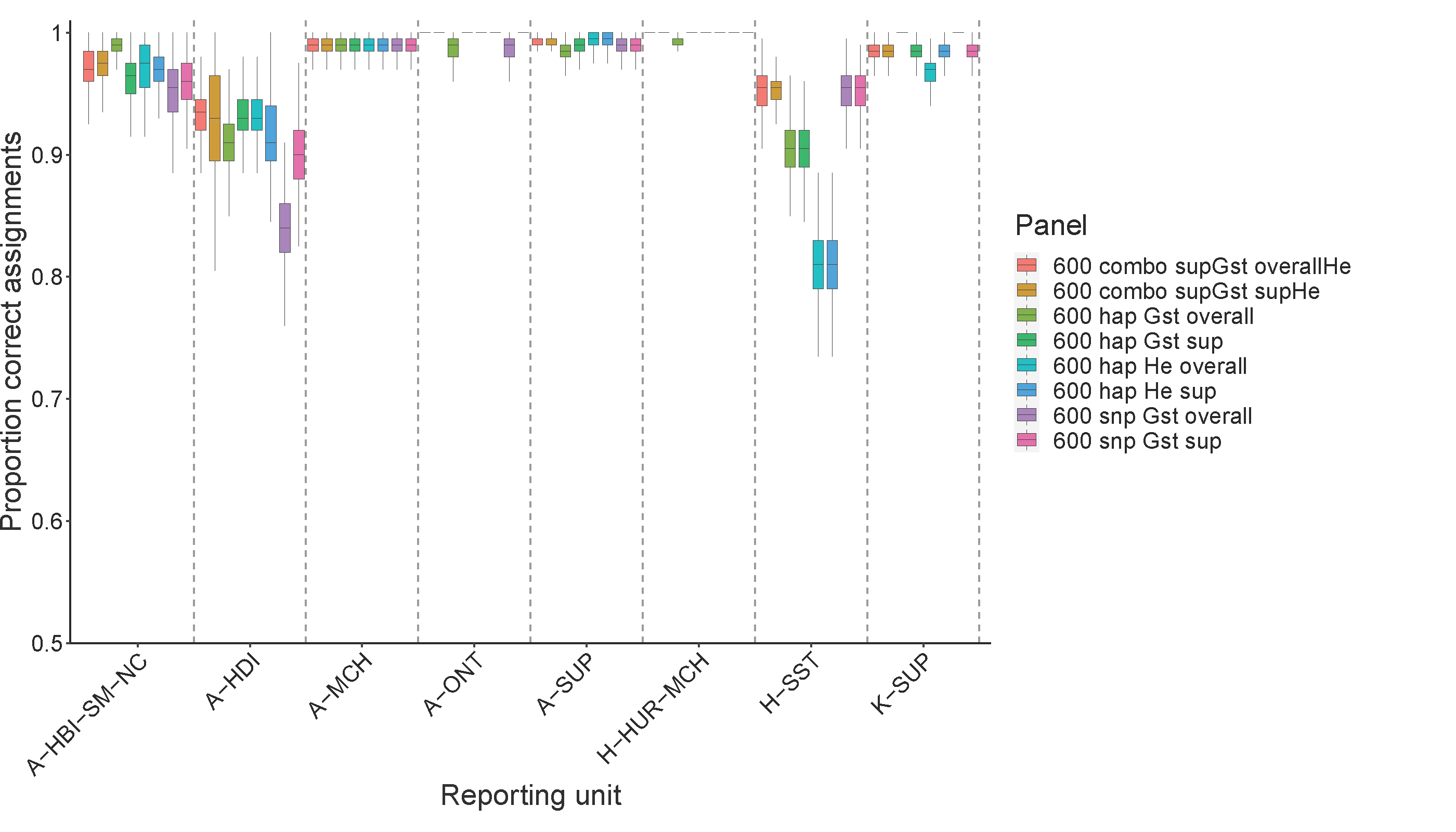

**Supplemental Figure 2**. Simulated performance of different combinations of loci for assignment of individuals to reporting units. Reporting units and locus combinations are described in Table 1 in the manuscript & and Supplement Table 1.

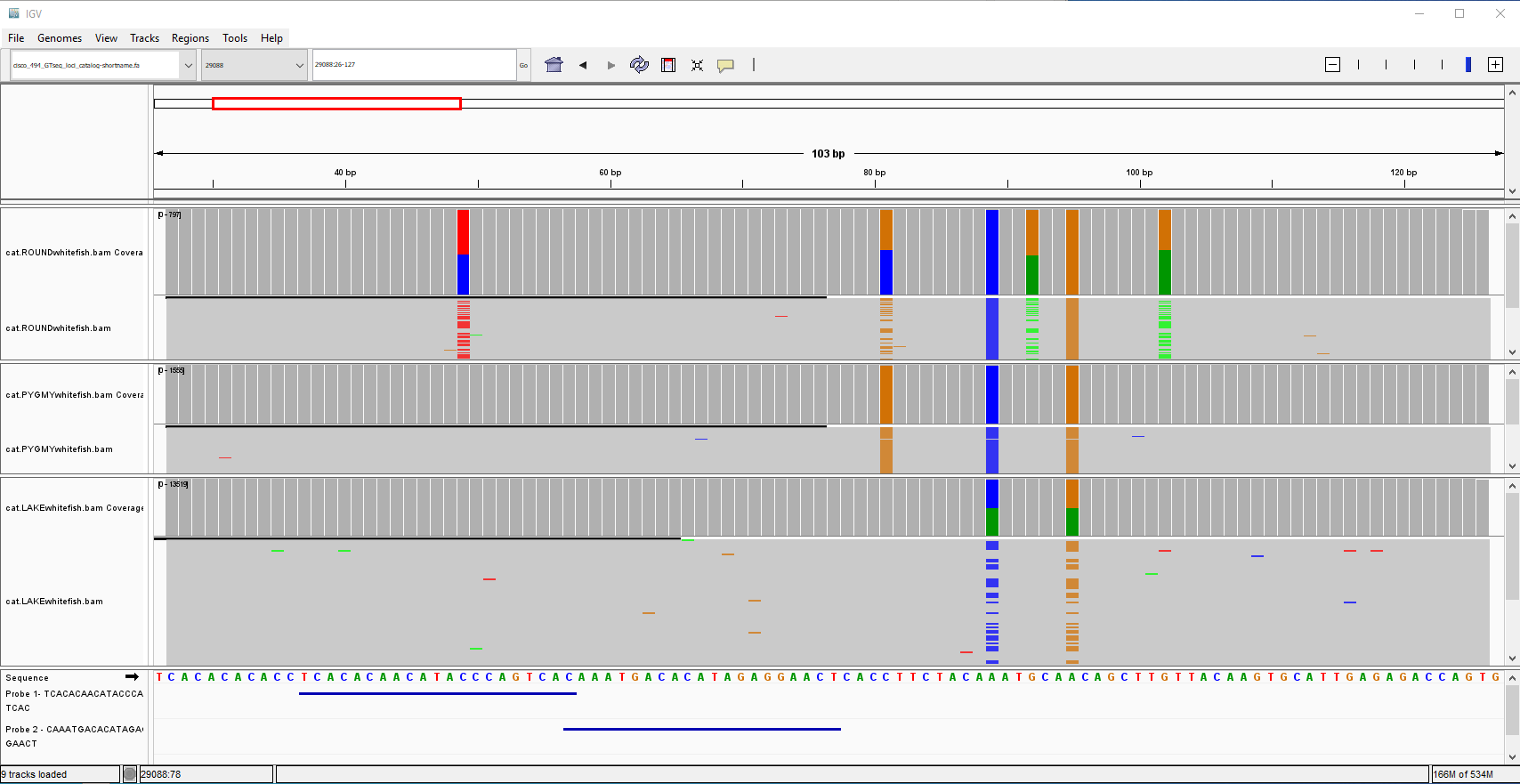

**A**

**B**

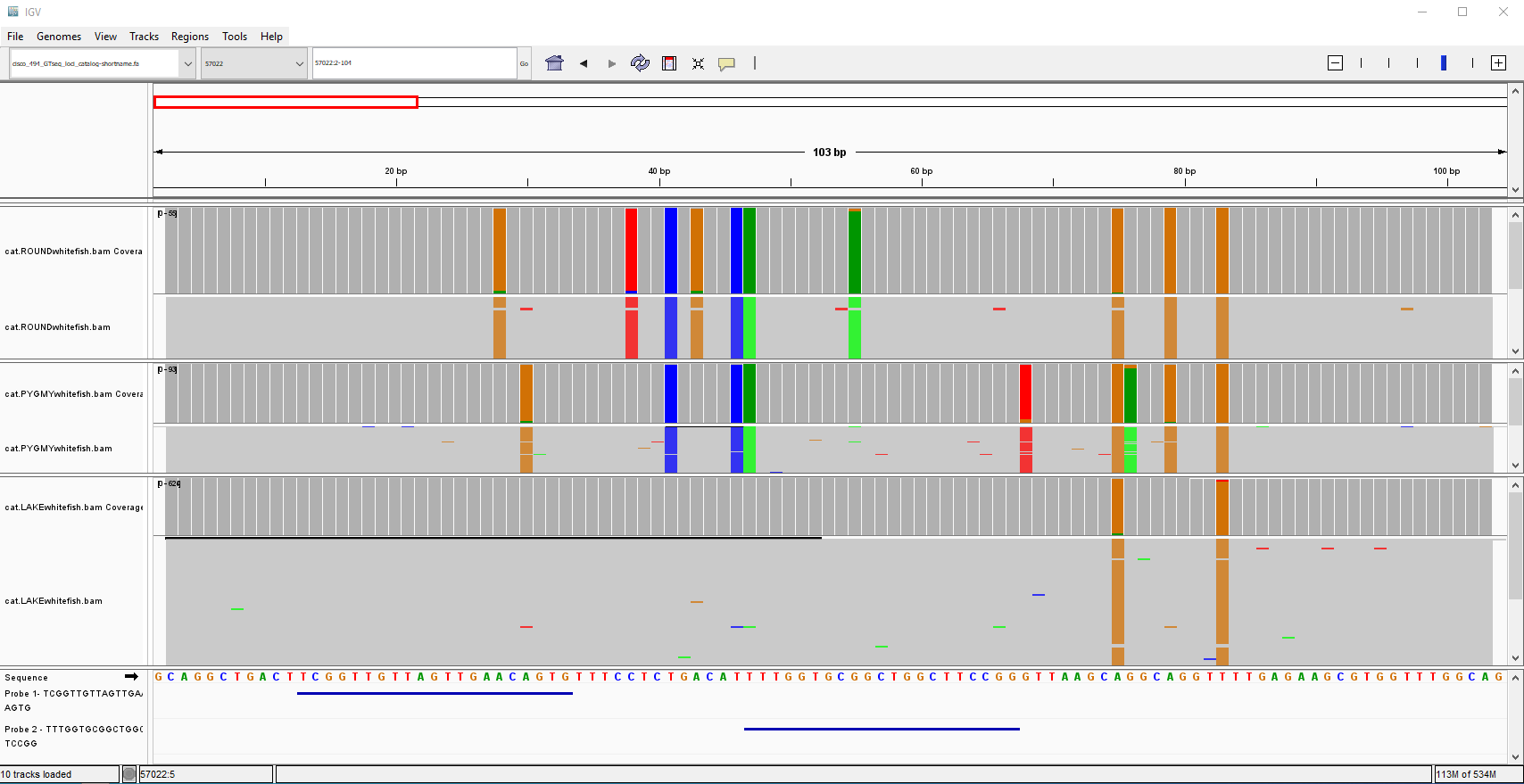

**Supplemental Figure 3.** Round Whitefish, Pygmy Whitefish, and Lake Whitefish BAM alignments in IGV for GT-seq panel locus 29088 (A) and 57022 (B). Panel A: the first probe (blue lines in bottom panel of window) for locus 29088 was redesigned to account for the variant SNP in Round Whitefish. Panel B: probes were not redesigned for locus 57022 because of variant complexity (multiple fixed SNPs in Round and Pygmy Whitefishes that would result in two or more degenerate sites in each probe).

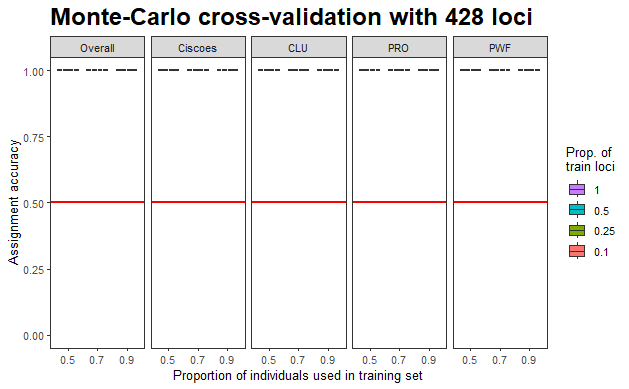
 **Supplemental Figure 4.** Assignment accuracy plot for Lake Whitefish (CLU), Round Whitefish (PRO) and Pygmy Whitefish (PWF) using a whitelist of 428 SNPs that were polymorphic in *Prosopium* whitefishes. All combinations of loci and individuals tested resulted in 100% assignment accuracy.

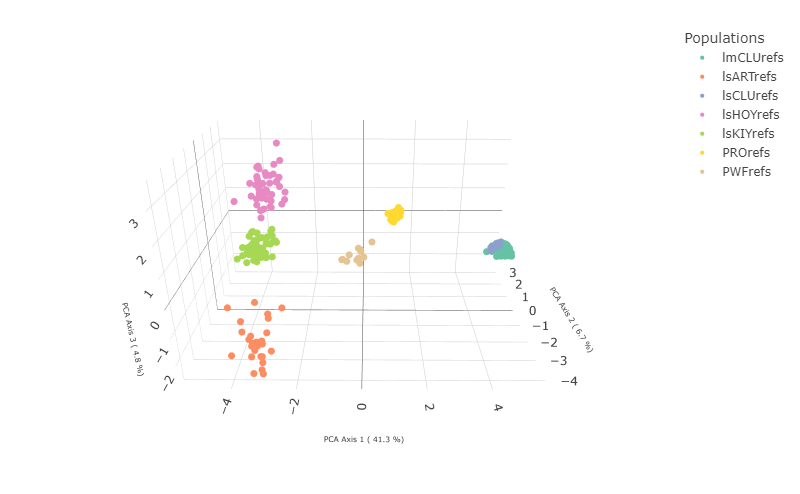

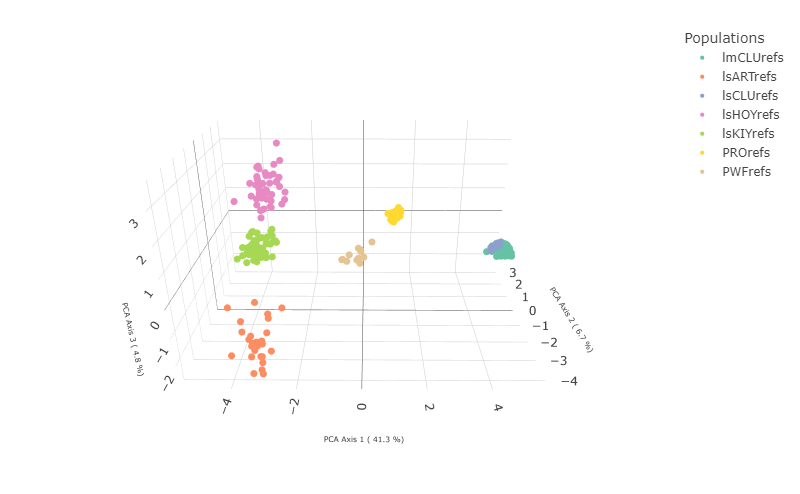

**Supplemental Figure 5.** Three-dimensional principal components analysis of reference individuals using a whitelist of 428 SNPs that were polymorphic in *Prosopium* whitefishes. lmCLUrefs – Lake Michigan Lake Whitefish, lsCLUrefs – Lake Superior Lake Whitefish, lsARTrefs – *C. artedi*, lsHOYrefs – *C. hoyi*, lsKIYrefs – *C. kiyi*, PROrefs – Round Whitefish, PWFrefs – Pygmy Whitefish.

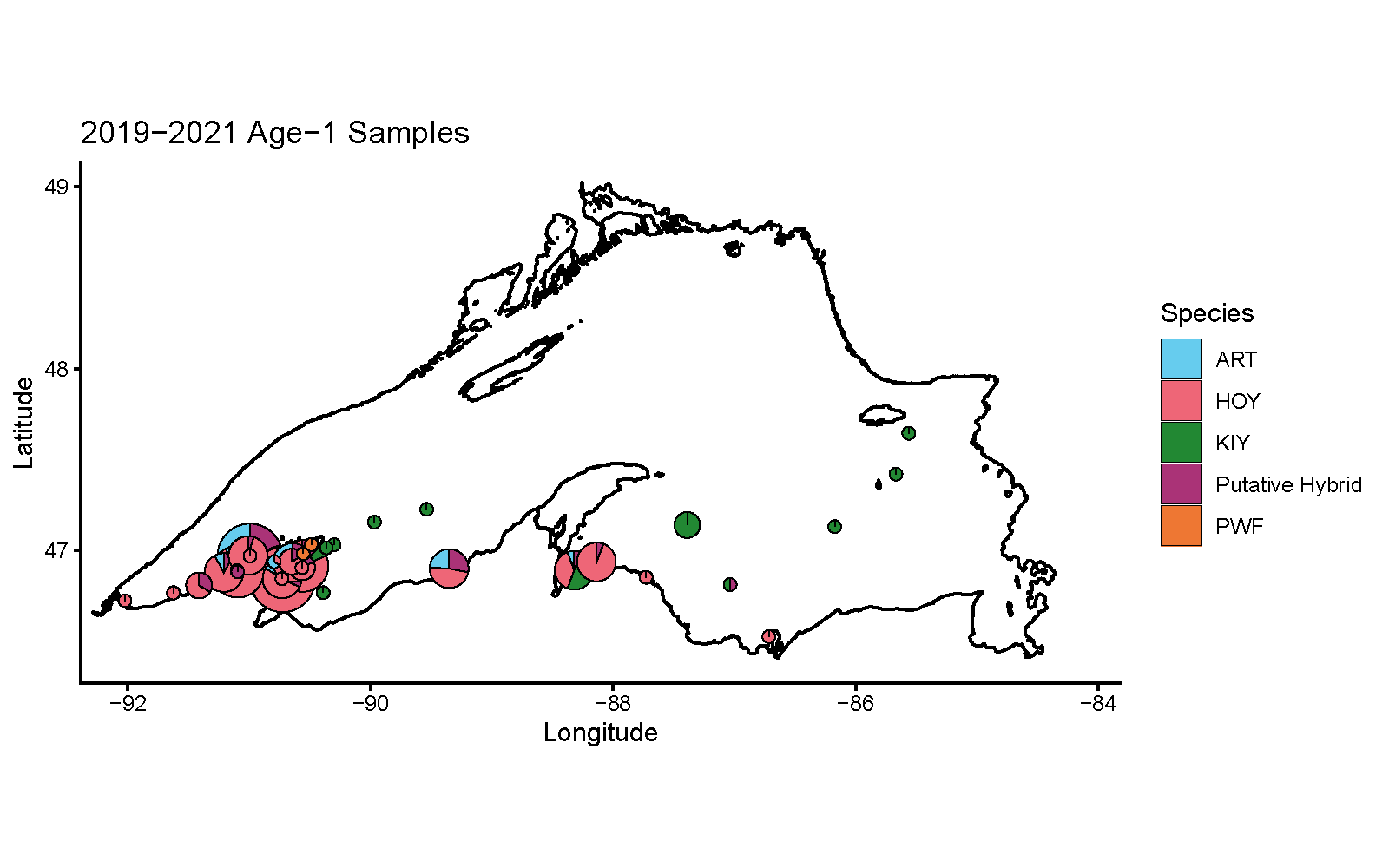

**Supplemental Figure 6.** Spatial distribution of genetically identified age-1s collected from Lake Superior in 2019-2021. Pie chart size is proximal to number of larvae from each collection site.
