## Supplemental File 2 GTseq Lib Protocol for "A genomic tool to tackle cryptic diversity demonstrates the potential for off-target use of GT-seq panels"

**SUPPLEMENTAL FILE 2, GT-seq panel library preparation protocol**

**A genomic tool to tackle cryptic diversity demonstrates the potential for off-target use of GT-seq panels**

Amanda S. Ackiss, Mark R. Vinson, Ann J. Ropp, Kristen M. Gruenthal, Trevor J. Krabbenhoft, Joseph V. Siegel, Wendylee Stott, Daniel L. Yule, Wesley A. Larson

.

Disclaimer: Any use of trade, firm, or product names is for descriptive purposes only and does not imply endorsement by the U.S. Government.

**SNPPCR**

Reaction mixture is 2 μL of DNA combined with 2 μL of 0.5 μM of the multiplexed primer pool and 6 μL Qiagen® Multiplex PCR Master Mix. Polymerase chain reaction (PCR) is run on a thermocycler with the following conditions: 5 minutes at 95°C for initial denaturation, 5 cycles of 95°C for 30 s, 57°C for 2 min with 5% or 0.3°C/min ramp, and 72°C for 30 s, followed by 10 cycles of 95°C for 30 s, 65°C for 30 s, and 72°C for 30 s, finishing with a 5 minute final elongation.

**BARCODING**

3 μL of SNPPCR product is diluted with 37 μL of molecular-grade water. 2 μL of the diluted PCR product is combined with 2 μL of 5 μM i05 index, 1 μL of 10 μM i07 index, and 5 μL Qiagen® Multiplex PCR Master Mix. Primers and custom indexes used are reported in Supplemental File 4. Indexing PCR is run on a thermocycler with the following conditions: 95°C for 15 minutes, 10 cycles of 95°C for 10 s, 65°C for 30 s, 72°C for 30 s, followed by a final elongation of 72°C for 5 minutes.

**NORMALIZATION**

All 10 μL of indexed library is transferred to a SequalPrep™ Normalization Plate (Applied Biosystems™) for normalization following manufacturer’s protocol.

**BEAD SIZE SELECTION**

Normalized libraries are then pooled and 200 μL of library pool is taken through a 0.6X-1.0X double-sided magnetic bead size selection with AMPure XP beads (Beckman Coulter). Final library product is eluted off beads with 35 μL Tris low-EDTA.

To ensure that a target product length of ~232bp bp was achieved, 2 μL of library is run on an Invitrogen E-Gel EX 2% gel and double-stranded DNA is quantified with a Qubit (ThermoFisher). Preparation of library pools from 95 samples and 1 no template control (negative) typically yields a concentration of 1-2 ng/μL.
